# Convergence of Heteromodal Lexical Retrieval in the Lateral Prefrontal Cortex

**DOI:** 10.1101/2020.11.30.405746

**Authors:** Alexander A. Aabedi, Sofia Kakaizada, Jacob S. Young, Olivia Wiese, Claudia Valdivia, Mitchel S. Berger, Daniel H. Weissman, David Brang, Shawn L. Hervey-Jumper

## Abstract

Lexical retrieval requires selecting and retrieving the most appropriate word from the lexicon to express a desired concept. Prior studies investigating the neuroanatomic underpinnings of lexical retrieval used lesion models that rely on stereotyped vascular distributions, functional neuroimaging methods that lack causal certainty, or awake brain mapping that is typically limited to narrow cortical exposures. Further, few studies have probed lexical retrieval with tasks other than picture naming and when non-picture naming lexical retrieval tasks have been applied, both convergent and divergent models emerged. Because of this existing controversy, we set out to test the hypothesis that cortical and subcortical brain regions specifically involved in lexical retrieval in response to visual and auditory stimuli represent overlapping neural systems. Fifty-three patients with dysnomic aphasia due to dominant-hemisphere brain tumors performed four language tasks: picture naming, auditory naming, text reading, and describing line drawings with correct syntax. A subset of participants also underwent the Quick Aphasia Battery which provides a validated measure of lexical retrieval via the word finding subtest. Generalized linear modeling and principal components analysis revealed multicollinearity between picture naming, auditory naming, and word finding, implying redundancies between the linguistic measures. Support vector regression lesion-symptom mapping across participants was used to model accuracies on each of the four language tasks. Picture naming and auditory naming survived cluster-level corrections. Specifically, lesions within overlapping clusters of 8,333 voxels and 21,512 voxels in the left lateral PFC were predictive of impaired picture naming and auditory naming, respectively. These data indicate a convergence of heteromodal lexical retrieval within the PFC.

**Importance of the Study:** Lexical retrieval (i.e., selecting and retrieving words to convey desired concepts) is a crucial component of language processing. However, existing studies of the neuroanatomic underpinnings of lexical retrieval lack causal relationships and have provided conflicting evidence, suggesting both convergent and divergent models. In order to resolve these conflicting models, we used lesion-symptom mapping to investigate lexical retrieval in 53 patients with dominant-hemisphere brain tumors. We observed significant associations between performance on visual and auditory naming tasks. Further, performance on these tasks predicted performance on a validated neuropsychological measure of lexical retrieval. Critically, multivariate, nonparametric lesion-symptom mapping within a brain tumor framework revealed that lesions in overlapping regions of the left lateral prefrontal cortex (PFC) predict impaired visual and auditory naming. In a clinical context, this approach to identifying causal brain-behavior relationships could help to guide brain tumor therapies such as cytoreductive surgery and supportive rehabilitation services.

## Introduction

Lexical retrieval, the process by which a word that best conveys a given concept is selected from the lexicon, is universally required for natural language and impaired in nearly all forms of aphasia^1,2^. Models of lexical retrieval during speech production describe two main processing steps. First, the meaning of a word is accessed (i.e., lexical semantics). Next, the sound code is accessed (i.e., lexical phonology)^3–8^. Incongruencies exist in current lexical retrieval models in part because the process was traditionally examined using only visual picture naming tasks^6,9^. Specifically, various authors have proposed that picture naming begins with visual object recognition and retrieval of nonverbal conceptual knowledge of the visual stimulus. Next, the meaning of the word is accessed (lexical semantics). Finally, the lexical phonological word form is retrieved^8^. A number of studies have since incorporated non-visual, auditory naming tasks to provide a more ecologically valid and comprehensive representation of lexical retrieval^10^. However, rather than generate a unified representation of lexical retrieval that encompasses distinct sensory modalities, studies employing visual and auditory naming tasks have presented conflicting models.

Convergent models of lexical retrieval support overlapping visual and auditory neural systems within the frontal and temporal lobes. For example, Hamberger *et al*. (2001) used direct electrical stimulation (DES) to show that both visual and auditory naming sites converge within the posterior temporal cortex^11^. Similarly, Hamberger *et al*. (2014) did not find any frontal or temporal regions unique to auditory naming via functional magnetic resonance imaging (fMRI)^12^.

Alternatively, divergent models support distinct anatomic correlates involved in lexical retrieval in response to visual and auditory stimuli. Malow *et al*. (1996) found “auditory only” naming sites primarily in posterior temporal regions using DES^13^. Hamberger *et al*. (2001)^11^ and Hamberger and Seidel (2009)^14^ reported that patients with lesions in the same area had impairments in visual but not auditory naming, and that the anterior temporal lobe conferred specificity for auditory naming.

These incongruent results may reflect limitations of the previous methodologies used to study heteromodal lexical retrieval. Indeed, DES is restricted to regions exposed during brain mapping surgery, may be affected by the administration of anesthetics, and induces nonphysiologic, backward propagation of action potentials, thereby limiting its spatial specificity^15–17^. Furthermore, studies employing DES often do not assess subcortical tissue, differentiate speech arrest (i.e., a transient dysfunction in general speech production) from true anomia/dysnomia, or match stimuli on content category. Finally, while functional imaging modalities such as electrocorticography, fMRI, and PET offer correlational insights with varying temporal and spatial precisions, they cannot generate conclusions about requisite brain areas with causal certainty^18^.

Lesion symptom mapping (LSM) is poised to identify cortical and subcortical regions that are necessary for a given task, either directly or indirectly via involvement of tissues connected to distant brain regions (diaschisis)^19^. Traditionally, LSM was implemented using a chronic stroke lesion model, thereby offering a view of language processing based on vascular territory. However, LSM has since been expanded to include lesions formed by intrinsic brain tumors, cytoreduction surgery, traumatic brain injury, and neurodegenerative disease^20–23^. Because dominant-hemisphere intrinsic brain tumors in particular lead to substantial rates of dysnomic aphasia^24^ and can encompass broad cortical and subcortical regions without confinement to stereotyped vascular distributions, they may serve as optimal lesion models to study lexical retrieval in response to visual and auditory stimuli.

In this study, we used a permutation-based, multivariate approach to LSM to identify regions necessary for heteromodal lexical retrieval. Because recent data using a combination of functional imaging and DES offer conflicting models of lexical retrieval, it remains unknown whether separate auditory- and visual-only regions exist, or whether the neural systems subserving lexical retrieval exist through distributed convergence zones^25^. Thus, our aims were twofold: 1) to determine whether lesions in frontotemporal regions selectively impair auditory versus picture naming and 2) to identify causal convergence zones using a lesion-symptom mapping framework.

## Materials and Methods

### Participants

Eighty-nine adult patients presenting with World Health Organization (WHO) II-IV gliomas and brain metastasis at the University of California San Francisco (UCSF) between 2017 and 2020 were recruited in a longitudinal prospective clinical trial of language and neurocognitive outcomes (NCI-2020-02286). This included fifty-three adult patients (participants) with dominant hemisphere tumors and a disease- and age-matched control cohort of thirty-six patients (controls) with non-dominant hemisphere tumors. All patients provided written informed consent for study enrollment in accordance with UCSF’s institutional review board. Hemisphere of language dominance was established via magnetoencephalography (MEG)^26^. Tumor histology was classified according to the WHO 2016 classification of CNS tumors^27^. Fresh tissue samples were imaged using stimulated Raman histology with cell counting of core specimens using previously established protocols^28^.

### Experimental Design and Statistical Analysis

#### Preoperative Language Assessments

Each participant and control underwent a language assessment one day prior to cytoreduction surgery using the validated Quick Aphasia Battery (QAB) which provides weighted scores for each of seven predefined language domains (“subtests”): word comprehension, sentence comprehension, word finding, grammatical construction, motor speech, repetition, and reading^29^. The QAB was used in this study given its prior use in adult brain tumor patients and ability to provide rapid (yet comprehensive) assessments of language functions. Furthermore, the QAB was particularly valuable for the purposes of this study as it provides a validated measure of lexical retrieval through its word finding subtest. Nine participants did not undergo QAB testing and were excluded from components of the analysis that required QAB scores. QAB scores between participants and controls were compared using the Wilcoxon rank-sum test after confirming non-normality of scoring distributions both visually and via the Shapiro-Wilk test. This comparison was 80% powered to identify a medium-to-large effect size (Cohen’s *d* of 0.67) using a significance level of 0.05. To identify sources of potential confounding in language scores, categorical comparisons of demographics and baseline clinical data between study participants and controls were performed with chi-squared tests.

In addition to completing the QAB, each of the fifty-three participants performed four additional language tasks for use in lesion-symptom mapping. These tasks consisted of naming pictorial representations of common objects and animals (Picture Naming, PN), reading two-syllable text (Text Reading, TR), naming common objects and animals via auditory descriptions (Auditory Naming, AN) and describing line drawings with intact syntax (Syntax, Syn)^17,30^. The correct answers for all four tasks were matched on word frequency (i.e. commonality within the English language) using SUBTLEX_WF_ scores provided by the Elixcon project and content category^31^ (http://elexicon.wustl.edu/).

All of the tasks were delivered on a laptop with a 15-inch monitor (60 Hz refresh rate) that was positioned two feet away from the seated patient in a quiet clinical setting. Task stimuli were randomized and presented using PsychToolbox^32,33^. Slides were manually advanced by the research coordinator either immediately after the participant provided a response or after six seconds if no response was given. All of the tasks were scored on a scale from 0 to 4 by a trained clinical research coordinator who was initially blinded to all clinical data (including imaging studies) using the guidelines provided by the QAB. No participants had uncorrectable visual or hearing loss.

To identify which language tasks were most strongly associated with the validated measure of lexical retrieval provided by the QAB (i.e., the word finding subtest), generalized linear models were fitted between the word finding subtest and each of the four language tasks. Principal components analysis (PCA) was performed to collapse the five language measures into a smaller set of dimensions (i.e. principal components) that account for a majority of the variance in the dataset. By removing redundancies in the behavioral data, PCA is able to identify common cognitive constructs. In this study, the largest resulting principal component (i.e., defined by a weighted combination of our five language measures) was used in lesion-symptom mapping to identify brain regions subserving a common cognitive construct. All statistical analyses were performed on R version 3.6.2. Corrections for multiple comparisons were made by controlling the family-wise error rate with the Holm-Bonferroni method^34^.

#### Magnetic resonance imaging and pre-processing

Each participant underwent a standard preoperative imaging protocol on a 3 T scanner including a) pre- and post-contrast T1-weighted imaging with gadolinium and b) T2-weighted fluid attenuation inversion recovery (FLAIR) imaging with slice thicknesses between 1 and 1.5 mm^30^. All imaging was performed within three days of the language assessments.

Lesions were segmented either manually or semi-automatically using ITK-SNAP 3.8.0 (http://www.itksnap.org/) by a trained co-author blinded to language outcomes scores (AAA)^35^. The borders of contrast enhancement on T1-weighted post-gadolinium sequences were used to identify lesion boundaries in patients with WHO IV contrast enhancing tumors (n = 43 patients). Abnormal FLAIR signal was used to identify lesion boundaries in patients with WHO II and III non-enhancing tumors (n = 10)^21,36^. Accuracy of the lesion masks was independently confirmed by a separate examiner also blinded to language outcomes scores (SHJ) and disagreements were resolved by reaching a consensus.

Using Clinical Toolbox on SPM12 (https://www.fil.ion.ucl.ac.uk/spm/software/spm12/), each anatomical T1-weighted image was normalized to the standardized Montreal Neurological Institute template (MNI152)^37,38^. The resulting transformation matrix was applied to the participant’s corresponding lesion mask to facilitate comparisons across participants (i.e., after placing all of the participants’ lesions into the same standardized space). For participants with non-enhancing lesions, the T2-weighted FLAIR derived lesion mask was registered to the anatomical image prior to normalization.

#### Lesion-Symptom and Statistical Analysis

Support vector regression lesion-symptom mapping (SVR-LSM) was performed to identify the precise anatomic locations associated with lower scores on each of the four language tasks (PN, TR, AN, and Syn) and the first principal component derived from PCA^39^. Analyses were conducted using the SVRLSMgui package (https://github.com/atdemarco/svrlsmgui/) for MATLAB which provides a permutation-based multivariate approach to lesion-symptom mapping with significant advantages over mass-univariate analyses^40^. As opposed to traditional voxel lesion-symptom mapping (VLSM), by including every lesion voxel as a simultaneous covariate, SVR-LSM accounts for the inherent spatial autocorrelation of brain tumors^41,42^. Additionally, by performing random permutations of the behavioral data to create null distributions, it is robust to the Type I and II errors that often arise from false discovery rate and Bonferroni methods for multiple comparisons corrections, respectively^43^.

Total lesion volume was linearly regressed out of both the lesion masks and language scores to limit the biasing impact of large lesions on language impairment while retaining sensitivity to fluctuations in task performance. For inclusion in each task-based analysis, each voxel was required to include at least two participants with overlapping lesions^44^. A threshold of p < 0.005 was used to determine statistical significance for individual voxels while a threshold of p < 0.05 was used for cluster-level (groups of contiguous voxels) corrections after performing 10,000 permutations of the behavioral data. Hyperparameters were left at their default settings (gamma of 5 and cost of 30). Analyses were fully parallelized on a high-performance computer with 32 cores at 4.2 GHz and 256 GB of random-access memory. SPM12 was used to generate statistical parametric maps and BrainNet Viewer (https://www.nitrc.org/projects/bnv/) to generate three-dimensional representations of the resulting statistical maps^45^.

## Results

Demographics and clinical data are summarized in Table 1. There were no significant differences in mean age, handedness, education, or oncologic features between participants and controls. Participants were significantly more likely to be male and to have left-sided tumors. Tissues sampled from within regions of FLAIR or T1 post gadolinium signal abnormality from three different participants with WHO grade II, III and IV gliomas are presented in Figure 1. All participants and controls had WHO II-IV gliomas as defined by WHO 2016 molecular sub classifications and imaged tissues demonstrate cellular neoplasms with extensive disruption of normal cytoarchitecture.

**Table 1:** Comparisons between categorical variables were made with the chi-squared test while continuous variables were compared with the Wilcoxon rank-sum test. Handedness was determined using preoperative magnetoencephalography. Education was self-reported in patients who completed the Neuro-QoL assessment. Pathologic diagnoses were made by board-certified pathologists using the World Health Organization (WHO) Revised Classification of Tumors in the CNS. Unk. – Unknown.

| Characteristic | Study Participants | Controls | p-value |
| --- | --- | --- | --- |
| *n* | 53 | 36 |  |
| Sex | 17 F, 36 M | 22 F, 14 M | 0.013 |
| Mean Age (SD) | 51.2 (17.3) | 49.4 (15.0) | 0.54 |
| Handedness | 3 L, 47 R, 3 unk. | 6 L, 27 R, 3 unk. | 0.19 |
| Education |  |  | 0.35 |
| High school | 10 | 6 |  |
| Some college | 11 | 7 |  |
| Bachelors | 11 | 13 |  |
| Graduate | 6 | 4 |  |
| Unk. | 15 | 6 |  |
| Tumor Laterality | 49 L, 4 R | 1 L, 35 R | $3.6 \times 10^{-16}$ |
| Tumor Grade/Pathology |  |  | 0.18 |
| WHO Grade I - II | 11 | 9 |  |
| WHO Grade III | 15 | 4 |  |
| WHO Grade IV | 24 | 23 |  |
| Metastasis | 3 | 0 |  |

**Figure 1:**
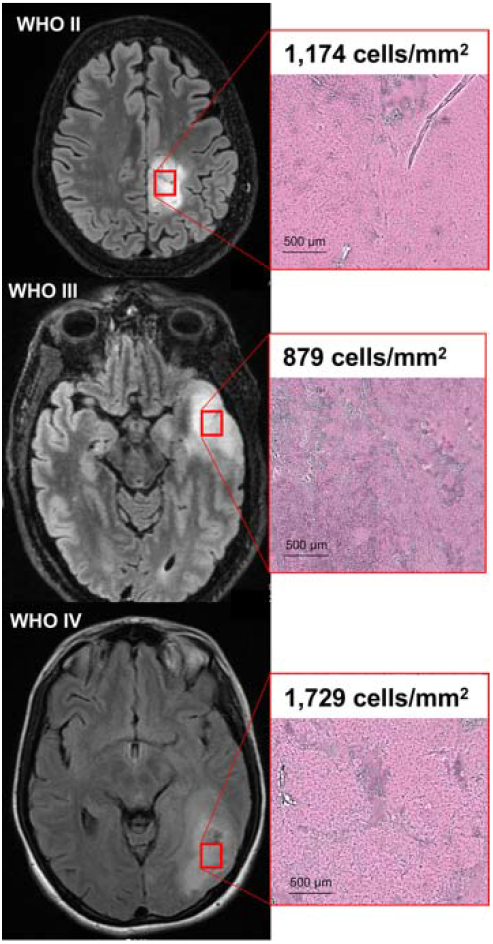
T2-weighted fluid attenuation inversion recovery (FLAIR) images of three study participants with World Health Organization (WHO) grade II, III, and IV glioma (left). Red boxes indicate regions of FLAIR signal abnormality biopsied for pathologic examination and cell-counting with Raman scattering microscopy (right). Pseudo-H&E images of fresh biopsy specimens demonstrate ablation of normal cytoarchitecture across all WHO tumor grades. Here, lesion cellularity and necrotic features escalate with increasing WHO grade.

Our main goal was to identify the neuroanatomical regions that are necessary for heteromodal lexical retrieval using a lesion-symptom model. As such, first, we compared language domain performance between participants and controls across all seven QAB subtests to establish whether or not participants had impairments in lexical retrieval. Compared to controls, participants exhibited significantly lower performance on word finding (p = 0.0096), but not on any of the six remaining subtests of the QAB (Fig. 2). Second, we sought to identify associations between each of the four language tasks (Table 2) and the word finding subtest using generalized linear models computed across all participants (Fig. 3A). Word finding was predicted by picture naming (*r* = 0.66, p = 0.000029) and auditory naming (*r* = 0.76, p = 0.0000001) but not by text reading (*r* = 0.37, p = 0.093) or Syn (*r* = 0.36, p = 0.093). Significant associations were also found between PN and AN (*r* = 0.80, p = 0.000000008), Syn and PN (*r* = 0.49, p = 0.0088), and Syn and AN (*r* = 0.68, p = 0.000016).

**Figure 2:**
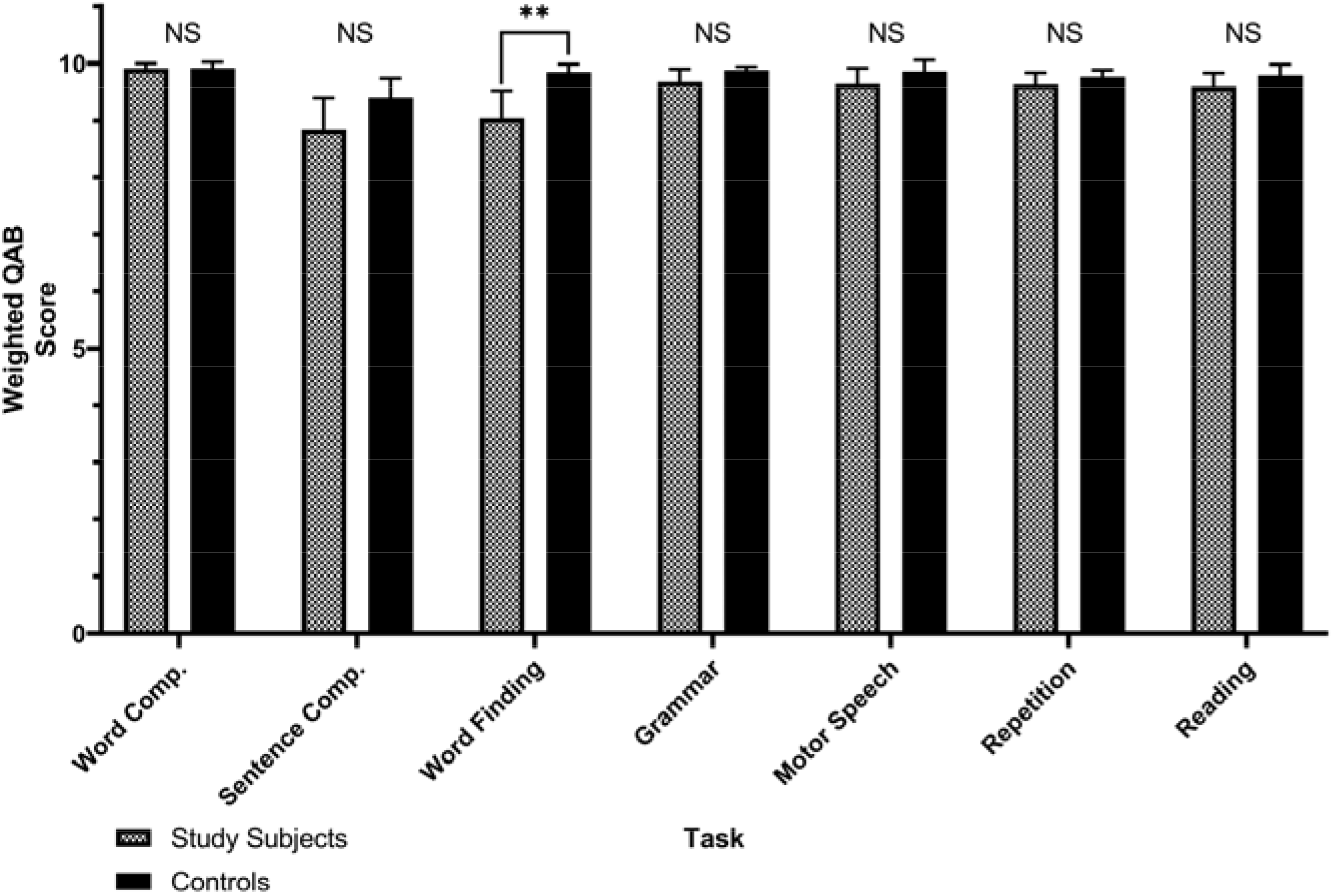
Pairwise Wilcoxon rank-sum tests between 44 patients with dominant-hemisphere intraparenchymal tumors (study participants) and 36 patients with non-dominant tumors (controls) on each of the seven predefined Quick Aphasia Battery (QAB) subtests. **p = 0.0096, corrected for multiple comparisons with the Holm-Bonferroni method. NS – not statistically significant (p > 0.05). Word Comp. – word comprehension. Sentence Comp. – sentence comprehension. Grammar – grammatical construction.

**Figure 3:**
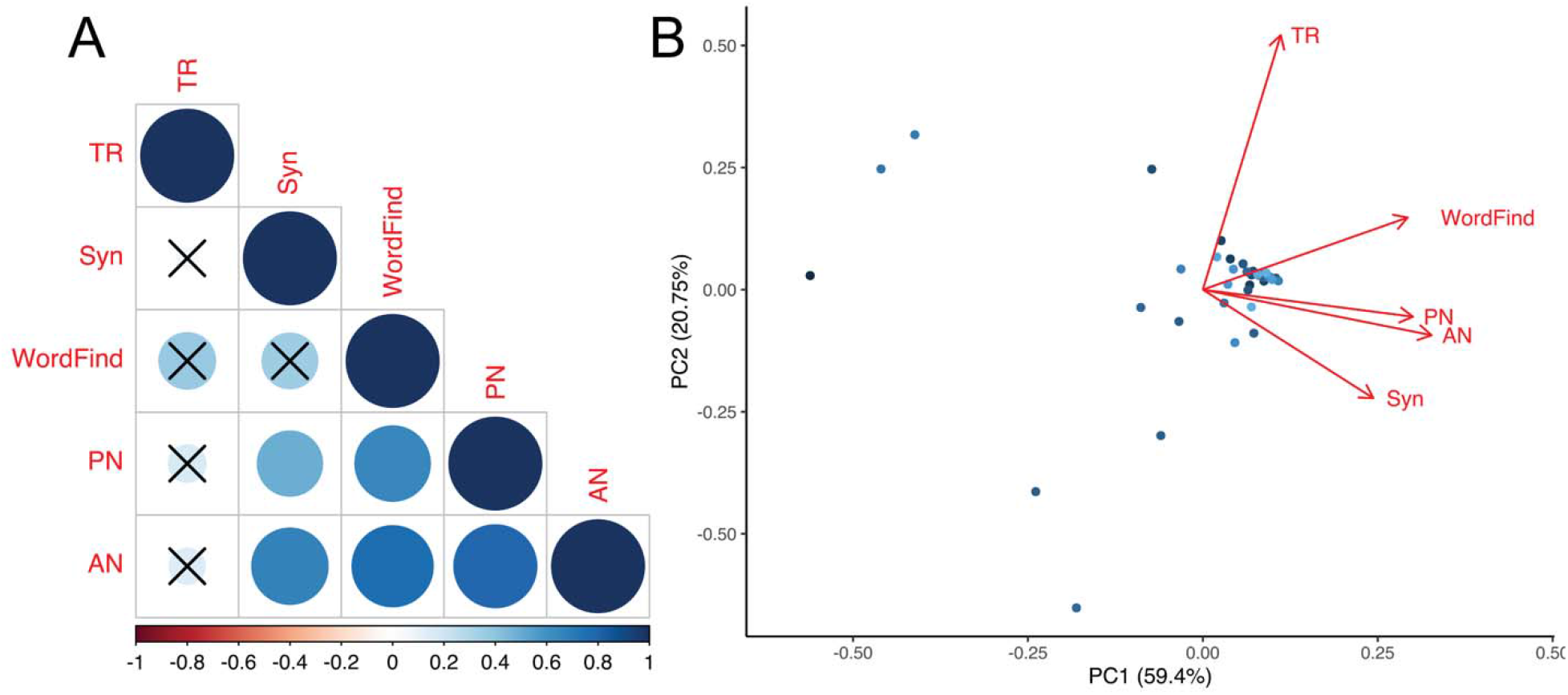
Operationalization of the word finding subtest of the QAB (WordFind) using four computerized language tasks: picture naming (PN), text reading (TR), auditory naming (AN), and syntax formation (Syn). ***A*,** correlogram summarizing the results of univariate generalized linear models fitted to each language task and WordFind. Color bar represents each correlation coefficient and crosses indicate non-significant associations (p > 0.05) after corrections for multiple comparisons. ***B*,** biplot between principal component 1 and 2 (PC1 and PC2) derived from principal components analysis (PCA) on WordFind and the four language tasks. Corresponding loadings are represented by the red arrows.

**Table 2:** Tasks were delivered in a quiet clinical setting and matched on word frequency. Each stimulus was scored from 0 – 4 using guidelines provided by the Quick Aphasia Battery.

| **Task (*n* trials)** | **Mean Score** | **Standard Error of Mean** |
| --- | --- | --- |
| Picture Naming (48) | 3.84 | 0.051 |
| Text Reading (27) | 3.97 | 0.014 |
| Auditory Naming (32) | 3.62 | 0.074 |
| Syntax (28) | 3.80 | 0.058 |

Given these results, we next performed PCA to a) reveal and remove redundancies in the behavioral data and b) identify a set of weighted combinations of the language measures that subsequently represent a single, common cognitive construct. PCA revealed two principal components (PC), PC1 and PC2, that were responsible for 59.4% and 20.75% of the variance in the five language measures, respectively (Fig. 3*B*). Picture naming, auditory naming, and word finding demonstrate the greatest collinearity with PC1. Text reading, on the other hand, is roughly collinear with PC2.

Having established the importance of picture naming and auditory naming in explaining the variance in the word finding subtest, we then used these two tasks in conjunction with SVR-LSM to uncover the anatomical regions necessary for visually- and auditorily-prompted lexical retrieval, respectively. Of the 53 participants, 48 had lesions with at least one overlapping voxel with another participant and were therefore included in the lesion-symptom mapping analysis. All five excluded participants had right-sided, dominant-hemispheric tumors (confirmed by MEG) with non-overlapping lesions. The final lesion overlap map used for SVR-LSM is outlined in green in Figure 4.

**Figure 4:**
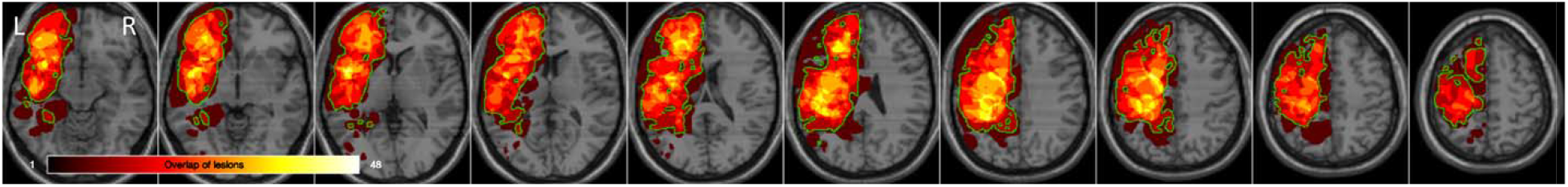
Map depicting regions in which at least two patients had overlapping lesions, project**ed** on the standard-space Montreal Neurological Institute template (MNI152), total *n* = 48. Regions in which actual overlap occurred and where the analysis was restricted to is outlined in green. Accordingly, five patients with non-overlapping right-sided tumors were excluded from lesionsymptom mapping.

Using this lesion map, SVR-LSM was then performed for each of the four language tasks testing the hypothesis that lesioned voxels are associated with lower task accuracies. For picture naming, one of eight clusters survived cluster-level corrections (8,333 voxels, p = 0.045). The resulting cluster for picture naming is centered in the left lateral PFC and encompasses Brodmann areas 10 and 45 – 48 (Fig. 5*A*). For auditory naming, one of ten clusters survived thresholding in an analogous region of the lateral PFC (Fig. 5*B*), except with greater involvement of subcortical areas (21,512 voxels, p = 0.0034). Notably, no significant clusters for auditory naming were observed in the temporal areas. The significant cluster in picture naming overlaps completely with, and accounts for 38.7% of, auditory naming’s significant cluster. Finally, SVR-LSM was performed using PC1 from the PCA. Only one of fourteen clusters survived cluster-level corrections (11,944 voxels, p = 0.019). This cluster overlaps with the resulting clusters from picture naming and auditory naming (Fig. 5*C*).

**Figure 5:**
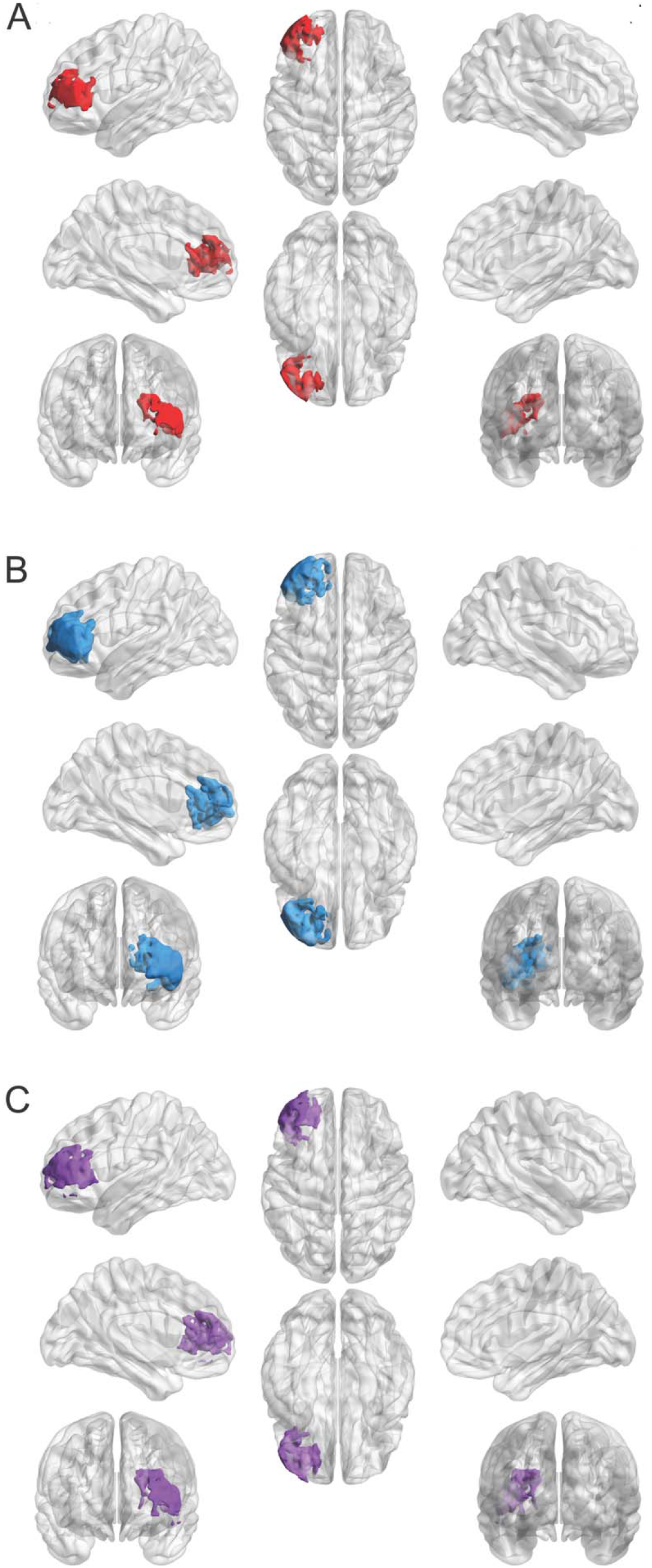
Results of support vector regression lesion-symptom mapping (SVR-LSM) after voxels thresholded at p < 0.005 underwent cluster-level corrections with 10,000 permutations (cluster threshold = p < 0.05). 3-Dimensional models of resulting clusters were generated using the Montreal Neurological Institute template via BrainNet Viewer (https://www.nitrc.org/projects/bnv/). ***A*,** Lesions in a cluster of 8,333 voxels in the left lateral PFC were predictive of impaired picture naming (p = 0.045). ***B*,** Lesions in a larger cluster of 21,512 were voxels predictive of impaired auditory naming (p = 0.0034). Clusters in *A* and *B* overlap completely with the cluster in *B* extending deeper into subcortical regions. ***C*,** Results of SVR-LSM using the principal component scores for each participant on the first principal component (PC1). This cluster consists of 11,944 voxels (p = 0.019) and demonstrates overlap with the clusters in *A* and *B*.

Neither TR nor Syn survived cluster-level corrections. For TR, no significant voxels were found after voxel-based thresholding. Syn, on the other hand, yielded smaller clusters of statistically significant voxels interspersed throughout the left anterior frontal lobe. However, the size of these clusters fell short of the critical threshold size that was derived from performing SVR-LSM on random permutations of the scores.

## Discussion

Cognitive models of lexical retrieval have been heavily influenced by visual picture naming tasks. Further, they implemented mixed methods that either lack causal certainty or precise spatial localization. Therefore, published results are mixed with respect to whether visual and auditory stimuli represent overlapping (convergent) or non-overlapping (divergent) neural systems. In this study, we propose a convergent model of lexical retrieval within a lesion-symptom framework for both visual and auditory inputs. First, we established that intrinsic brain tumors lead to extensive disruption of normal cytoarchitecture (Fig. 1), leading to selective impairments in word finding (Fig. 2). This finding is in line with previous reports of the prevalence of selective dysnomia in patients with dominant hemisphere intrinsic brain tumors^46^. Furthermore, it supports the use of our brain tumor lesion model in particular to study lexical retrieval, since participants in other clinical populations tend to have additional confounding language impairments^47,48^.

Next, we demonstrated strong correlations between accuracy on picture naming and auditory naming, providing initial evidence of the existence of lexical retrieval pathways that are agnostic to the sensory modality of the input (Fig. 3). From a behavioral perspective, this finding has been replicated in lesion studies across several clinical populations. For instance, Hamberger and Seidel (2003) found that patients with left temporal lobe epilepsy had co-occurring impairments in visual and auditory naming compared to a) healthy controls and b) patients with right temporal lobe epilepsy^49^. Miller *et al*. (2010) and Hirsch *et al*. (2016) also reported significant associations between picture naming and auditory naming in lesion studies of patients with dementia^50,51^. Hirsch *et al*. in particular used PCA in a cohort of 458 patients to reveal a unique redundancy between picture naming and auditory naming that was not shared with any of their other twenty-five cognitive and linguistic measures. In the present study, PCA led to analogous results: the similarity in loadings between PN, AN, and word finding argue for a single linguistic construct (i.e. lexical retrieval) that can be differentiated from constructs contributing to other linguistic measures such as TR and Syn.

As predicted by the results of our language assessments, multivariate lesion-symptom mapping revealed overlapping clusters for picture naming, auditory naming, and PC1 in the lateral prefrontal cortex (Fig 5*A-C*). Notably, in contrast to prior DES studies^11,52^, a separate cluster specific to auditory naming was not identified in the anterior temporal lobe, a well-represented region in our lesion overlap mask. Our results instead align with a number of more recent, mixed-method studies that argue for a convergence hub in the lateral PFC for both visual and auditory inputs^25,53^. Specifically, by timing the evolution of cortical responses to the onset of task stimuli using electrocorticography, Forseth *et al*. (2018) showed that the inferior frontal gyrus (a component of the lateral PFC) serves as an interface between lexical and phonological pathways during both picture and auditory naming^25^. Taken with the results of the present lesion study, these findings provide compelling evidence that lexical retrieval indeed represents a unified cognitive construct that is agnostic to the input modality.

Notably, case reports and small clinical series of patients with modality-specific naming impairments such as visual (i.e. optic aphasia), auditory, and tactical anomia from brain lesions may seem to argue against the existence of a single construct subserving lexical retrieval^54–56^. However, these studies have not been widely validated in large clinical cohorts using rigorous image-based methods and thus cannot be adequately evaluated for confounding variables. Specifically, concurrent damage to primary sensory processing or association areas cannot be ruled out without the granularity provided by voxel-based analyses. For instance, Hamberger and Seidel (2009) reported that a cohort of fourteen patients with anterior temporal lobe lesions (defined as < 5 cm from the temporal pole) had impaired auditory, but not picture naming^14^. However, of those fourteen patients, only five had structural lesions. The remaining “anterior lesion” patients were defined by the presence of seizure foci on subdural or scalp EEG. Given a) the absence of lesion overlap maps for examination and b) the ability of epileptic foci to transiently impair remote brain areas, we cannot rule out the possibility that the observed differences in auditory and picture naming performance were due to confounding damage to auditory sensory processing areas (i.e. A1-A3). Indeed, the “auditory-only” naming sites identified by Malow *et al*. (1996) via DES (i.e. transient induction of iatrogenic lesions) are chiefly located in the superior temporal gyrus where various features of auditory stimuli are encoded^57–59^. Analogous concerns have also been raised about other modality-specific anomia syndromes^60^.

Although we did not observe modality-specific anomia in the present study, we acknowledge that accuracies on picture naming and auditory naming tasks were not perfectly concordant across participants. Indeed, while clusters for PN and AN overlap completely, the latter is considerably larger than the former. However, this result has been replicated in previous studies, and is thought to reflect increased competition between alternative words during auditory as opposed to picture naming^12,61^. The underlying etiology of this phenomenon is poorly understood and warrants future investigation.

### Limitations of the Present Study

It is first worth discussing the typical limitations that apply to any lesion-symptom analysis, namely that conclusions about behavioral contributions of brain areas lying outside our lesion masks cannot be made. For this reason, we expanded our analysis to include voxels in which at least two patients overlapped, rather than more restrictive cutoffs of five or more found in previous studies. By performing multivariate comparisons on an HPC that can support these increasingly memory- and computationally intensive analyses, we were able to avoid the reductions in statistical power that typically constrain studies employing mass-univariate tests with post-hoc corrections for multiple comparisons. While lowering the lesion overlap threshold may increase exposure to outlier effects, our implementation of nonparametric statistics for lesion-symptom mapping mitigate this concern by calculating exact p-values without any *a priori* assumptions about the underlying distribution. In our study, as in others, the impact of outliers was greatly attenuated by performing 10,000 random permutations of the input behavioral data to generate a null distribution for each test statistic^62^. Despite these measures, however, our lesion mask was not able to capture posteroinferior regions of the perisylvian network, thereby restricting our conclusions to the frontal and anterior temporal cortices and angular gyrus.

Furthermore, while there is some debate on whether brain tumors can serve as reliable clinical models for lesion-symptom analyses, this position can be extended to stroke for which VLSM was developed and is most commonly implemented. Because perfect clinical models for brain lesions likely do not exist, the literature may be best served by an increase, rather than a decrease, in the diversity of the clinical populations under study^63^.

## Conclusion

To summarize generalized linear modeling and principal components analysis revealed multicollinearity between picture naming, auditory naming, and word finding tasks. Support vector regression lesion-symptom mapping across participants was used to uncover associations between lower task accuracies on each of the four language tasks and lesioned voxels. Picture naming and auditory naming lesions demonstrated overlapping clusters within the left lateral PFC. This study demonstrates that cortical and subcortical brain regions involved in lexical retrieval from visual and auditory stimuli represent overlapping neural systems.

## Acknowledgements

We deeply appreciate insights from Sofia Jimenez and Matthew Soleiman within their respective fields of psycholinguistics and cognitive neuroscience.

